# phylopath: Easy phylogenetic path analysis in R

**DOI:** 10.1101/212068

**Authors:** Wouter van der Bijl

## Abstract

1. Confirmatory path analysis allows researchers to evaluate and compare causal models using observational data. This tool has great value for comparative biologists since they are often unable to gather experimental data on macro-evolutionary hypotheses, but is cumbersome and error-prone to perform.
2. I introduce `phylopath`, an `R` package that implements phylogenetic path analysis (PPA) as described by Von Hardenberg & Gonzalez-Voyer (2013). In addition to the published method, I provide support for the inclusion of binary variables.
3. I illustrate PPA and `phylopath` by recreating part of a study on the relationship between brain size and vulnerability to extinction.
4. The package aims to make the analysis straight-forward, providing convenience functions and several plotting methods, which I hope will encourage the spread of the method.
5. `phylopath` is released under the GPL-3 license, and is freely available on CRAN (**https://cran.r-project.org/web/packages/phylopath/index.html**) and GitHub (**https://github.com/Ax3man/phylopath**).

## Introduction

The comparative method is a critical tool to answer macro-evolutionary questions, and has been since the start of evolutionary biology itself. It is often the only way to assess the generality of evolutionary patterns. A drawback of the method is that it is observational, not experimental, and is therefore often said to be unable to evaluate causal mechanisms (Martins 2000). However, causal models *do* predict correlations between certain variables to exist, and other correlations to be absent. It is these predictions that are leveraged in path analysis (Shipley 2000a), a specific form of structural equation modeling, that uses regression to test these predictions. Specifically, as it is used here, we can define statements about which variables a causal model predicts to be independent, given certain co-variates, and test those independencies. If we find that they are not independent, i.e. we find a regression coefficient significantly different from zero, this can be interpreted as evidence against the causal model.

Consider a minimal example, where A causes B and B causes C, i.e. A → B → C. Since there is no direct causal link between A and C, only through B, this causal model predicts that A and C are independent, given B. This prediction can be tested with the regression model C ~ B + A, where the coefficient of A is predicted to be close to zero. In other words, we expect no effect of A that is additional to the effect of B, since all causal effects of A on C should be mediated by B. This rationale can be expanded to more complicated scenarios, and allow us to critically assess whether data supports a causal model. Similarly, several competing causal models can be compared, where we can assess which one is best supported by the data (Shipley 2000b, 2013; von Hardenberg and Gonzalez-Voyer 2013). Path analysis is of great potential value to comparative biologists since it allows for better use of observational data, and emphasizes a quantitative comparison of competing causal evolutionary hypotheses.

In comparative biology normal regression models cannot be used for path analysis since the assumption of independence of observations is violated, as closely related species are expected to be more similar (Felsenstein 1985; Pagel and Harvey 1991). This similarity by descent can be corrected for with phylogenetic comparative methods, and regression analysis can be performed using phylogenetic generalized least-squares (PGLS) models. Von Hardenberg & Gonzalez-Voyer (2013) showed that PGLS can be successfully employed to perform confirmatory path analysis, based on the d-separation method by Shipley (2000b), and termed it phylogenetic path analysis (PPA).

By its nature, PPA is complicated, time consuming and error prone. For the worked exercise in the book chapter outlining the method (Gonzalez-Voyer and von Hardenberg 2014), the reader needs to define a list of 46 total d-separation statements and fit 21 PGLS models, and then compile the results afterwards. This takes a lot of time, requires a lot of code, and the number of steps required increases the chance for errors. Moreover, manual procedures such as intermediary rounding of results can in some cases significantly alter the final results. Therefore, I hope that a specialized software implementation will greatly increase the reproducibility of the method, decrease research effort to perform the analysis and encourage the spread of the method by decreasing entry barriers.

## A worked example

### Dataset

I will illustrate the use of the package by recreating a small part of the analysis by Gonzalez-Voyer *et al.* (2016). This study focused on the possible influence of brain size on the vulnerability to extinction in 474 mammalian species. Note that the goal of the analysis presented here is merely instructional, the original paper present a much more thorough analysis and should be used for biological inference.

The data used in the study is included in the package as red_list and red_list_tree. The data includes seven variables, listed in table 1. Note that the species names are set as rownames and that these names match the tip labels of red_list_tree. This is how the package matches the observations to the phylogeny. In contrast to many other phylogenetic packages, it is not necessary to remove all species with missing values or to trim the tree. As long as all species with complete data occur in the tree the package will take care of the rest, and the user receives messages about removed species and trimming of the tree, based on those variables that are included in the causal model set.

**Table 1:** The seven variables used in the analysis.

| Variable | Description |
| --- | --- |
| Br | Brain size |
| B | Body size |
| P | Population density |
| L | Litter size |
| G | Gestation period |
| W | Weaning age |
| Status | Vulnerability to extinction, as Red list status |

### Defining the causal model set

We start out by defining various relationships common to all causal models. We assume that brain size is caused by body size (a result of allometry), gestation length is a causal parent of both litter size and weening age, and that body size is a causal parent of population density, since these are all well-established relationships in the literature. We want to control for allometric effects of body size, and therefore include a direct effect of body size on status and an indirect effect through litter size. Additionally we also assume that the population density and life history variables all affect the vulnerability to extinction (which I will refer to as *status*), to limit the number of models that needs testing.

Since we are interested in testing for direct and indirect effects of brain size, we will vary those effects. Following the original authors, when considering indirect effects, brain size is a causal parent of litter size, gestation period and weaning age. When looking at direct effects, brain size is directly causally linked to status. This then leaves us with four causal hypotheses: a null model where brain size is irrelevant, a model with a direct effect, a model with indirect effects and a model with both.

We can define these models using the `define_model_set()` function. We supply a list of formulas for each model, using `c()`. Formulas should be of the form child `~ parent`, or you can read the `~ as` “caused by”, and describe each path in your model. Multiple children of a single parent can be combined into a single formula: `child ~ parent1 + parent2`. The paths that are shared between all models, can be included using the `.common` parameter. So we define our four models:

~~~
**library**(phylopath)
m <- **define_model_set**(
  null = **c**(),
  direct = **c**(Status~Br),
  indirect = **c**(L~Br, G~Br, W~Br),
  both = **c**(Status~Br, L~Br, G~Br, W~Br),
  .common = **c**(Br~B, P~B, L~B+G, W~G, Status~P+L+G+W+B)
)
~~~

It is easy to forget a path, or to make a typo. It is therefore good to make a quick plot to check. You can either plot a single model with e.g. `plot(m$direct)`, or plot all of them at once (figure 1a):

**Figure 1:**
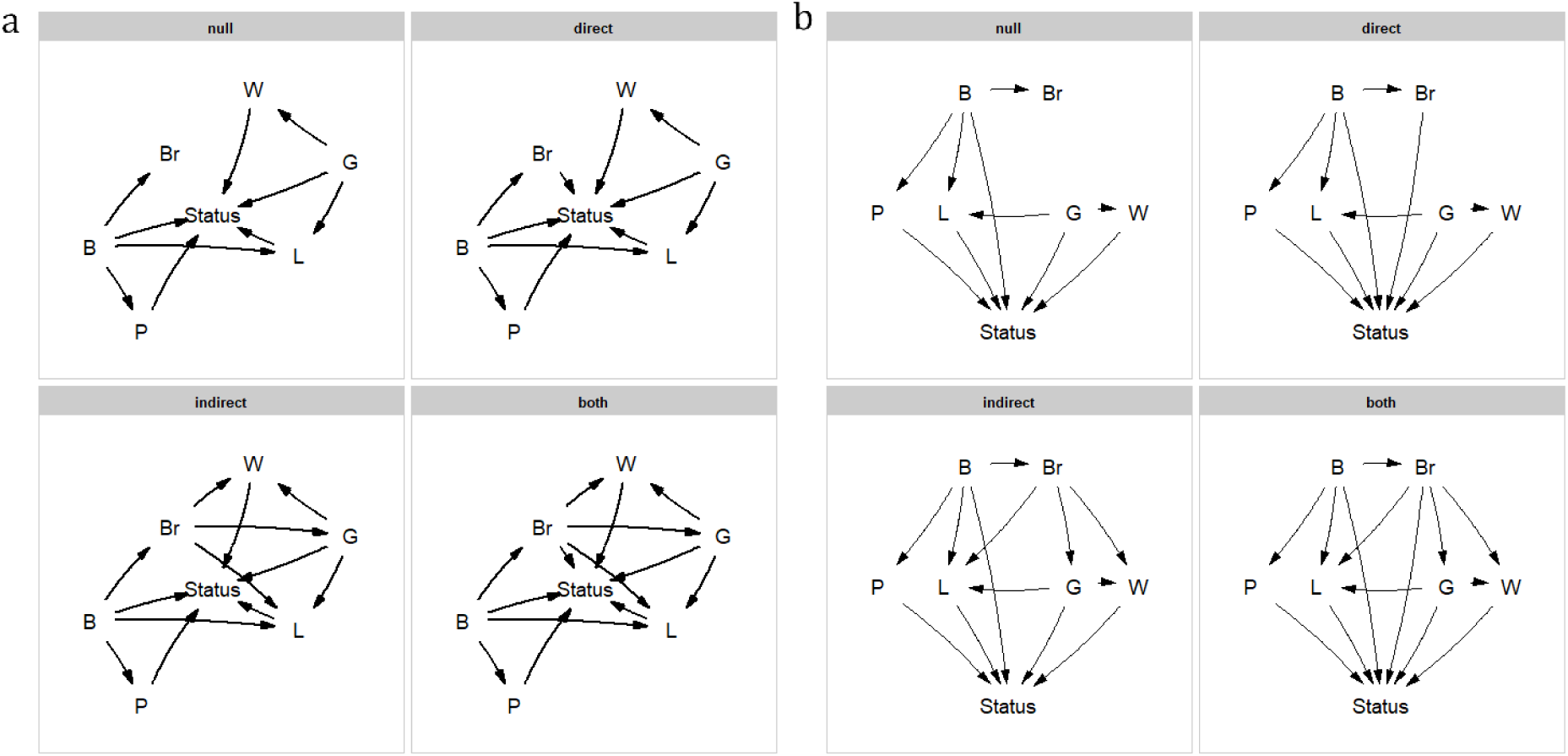
The model set, laid out algorithmically (*a*) and manually (*b*).

~~~
**plot_model_set**(m)
~~~

The nodes are laid out algorithmically. We can mimic the lay-out used in the paper by manually defining the coordinates in a data.frame (figure 1b), which in this case looks much better:

~~~
positions <- **data.frame**(
  name = **c**('B', 'Br', 'P', 'L', 'G', 'W', 'Status'),
  x = **c**(2:3, c(1, 1.75, 3.25, 4), 2.5),
  y = **c**(3, 3, 2, 2, 2, 2, 1)
)
**plot_model_set**(m, manual_layout = positions, edge_width = 0.5)
~~~

Defining your model set is perhaps the most crucial part of PPA. Since the method is confirmative and not explorative, you want to strike a good balance between complexity and interpretability.

### Evaluation of the hypotheses

~~~
p <- **phylo_path**(m, red_list, red_list_tree)
~~~

Printing the result gives us some basic information:

~~~
p
## A phylogenetic path analysis, on the variables:
## Continuous: G W B L P Status Br
## Binary:
##
## Evaluated for these models: null direct indirect both
##
## Containing 36 phylogenetic regressions, of which 18 unique
~~~

More importantly, asking for its summary and plotting it (figure 2) gives us the actual result of our comparison:

**Figure 2:**
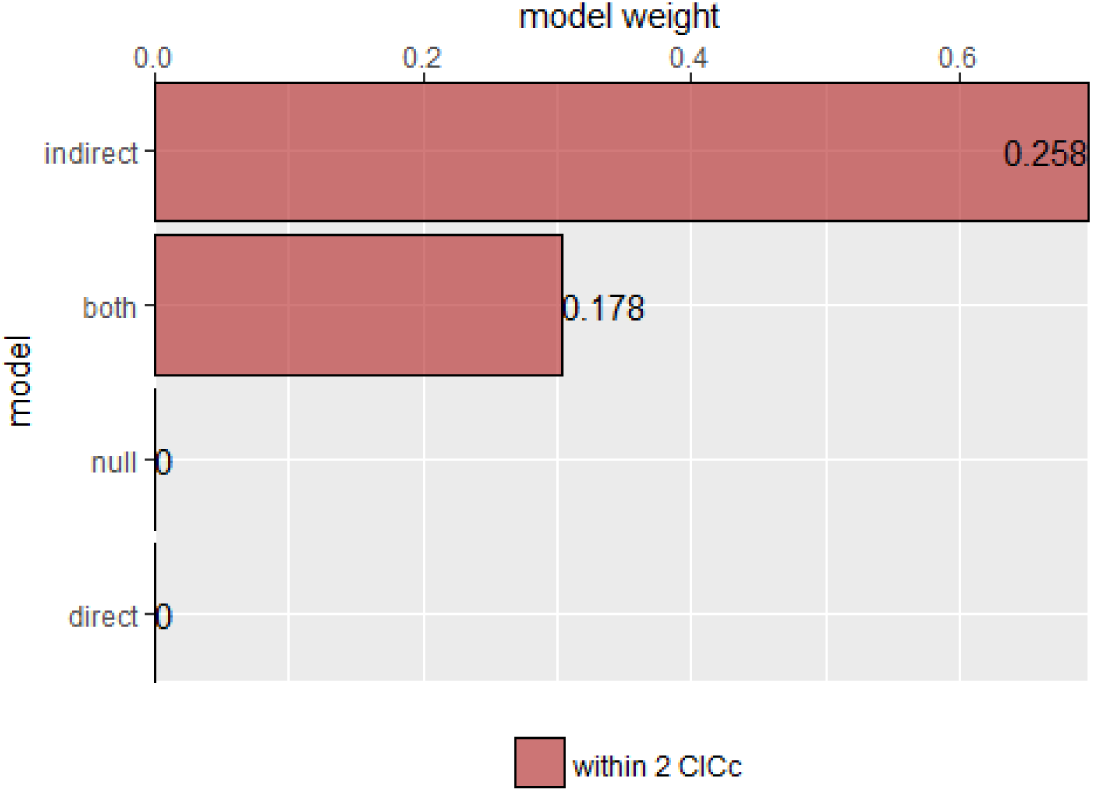
The relative importance of the four causal models.

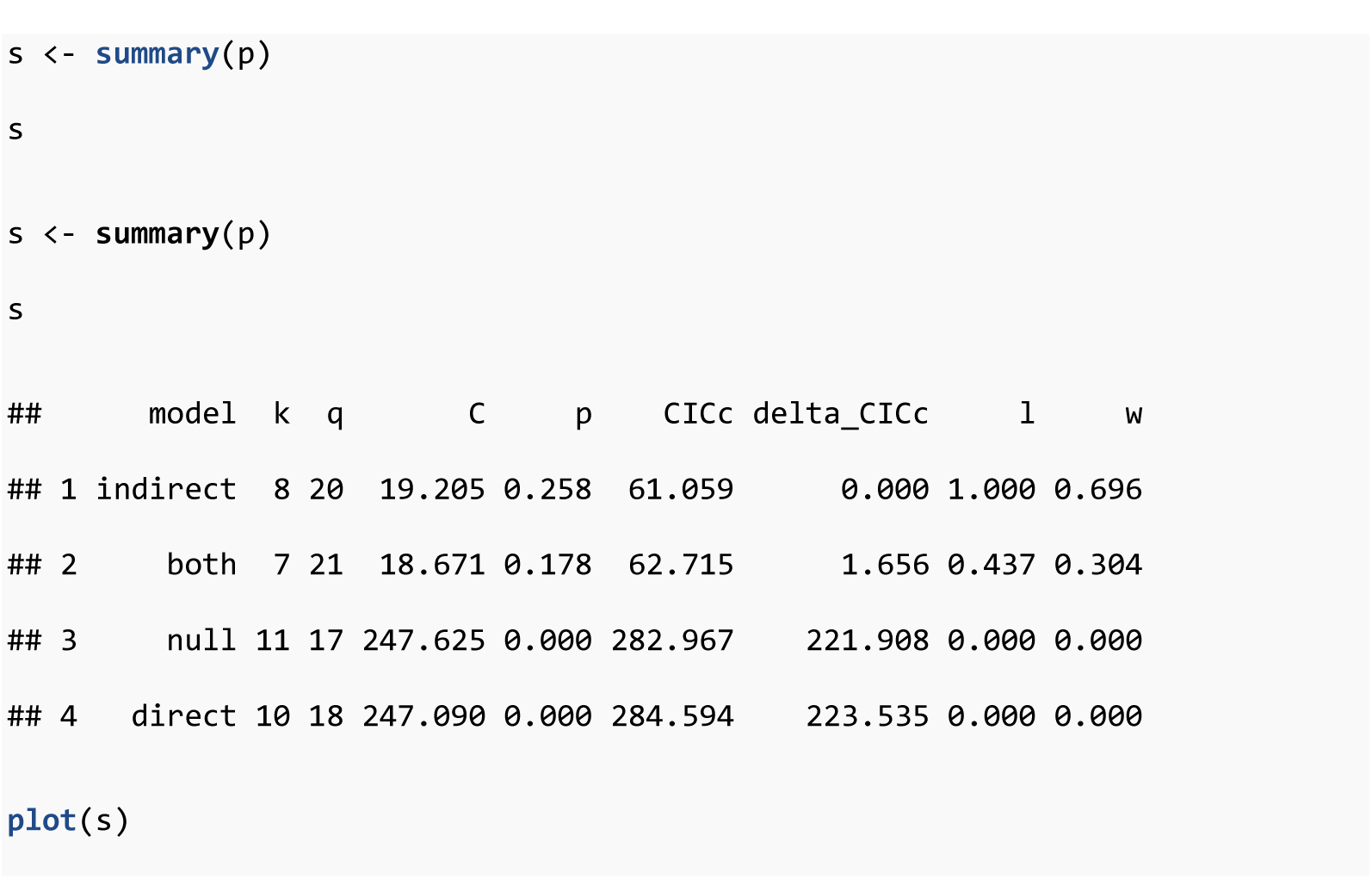

We can see that there is strong support for the indirect pathway. The addition of the direct path in the `both` model did lead to a small improvement (the C-statistic is lower) but not enough to put it ahead of the `indirect` model.

### Choosing a final model

So what is our best causal model? Firstly, the null and direct models are not supported since they have significant p-values, and should therefore be discarded. The indirect pathway is certainly important, but what about the direct pathway? There are several philosophies of dealing with this issue. In this particular case the two top-ranked causal models are directly nested, they share all the same paths except for one. We can think of this like nested regression models. We can say that the extra path should lower the CICc by at least some margin, often 2. In this case it does not, and we may elect to choose the top ranked model (see Arnold 2010 for a discussion on AIC and uninformative parameters).

After we have found our final model, we can estimate the relative importance of each of the paths. To estimate the paths in the highest ranked model, use the `best` function:

~~~
b <- **best**(p)
**plot**(b, manual_layout = positions)
~~~

This will return both the standardized regression coefficients, as well as their standard errors. The resulting plot is shown in figure 3. In order to get confidence intervals as well, you need to take bootstrap replicates using the boot argument: e.g. `b_ci <- best(p, boot = 500)`, which uses the bootstrap methods of the phylolm package (see “implementation notes”). This is disabled by default because it is slow. Using plot will give a visualization of the causal model. You can fit any arbitrary causal model that you evaluated with `choice`, so in this case `choice(p, “both”)` would give us the second ranked model.

**Figure 3:**
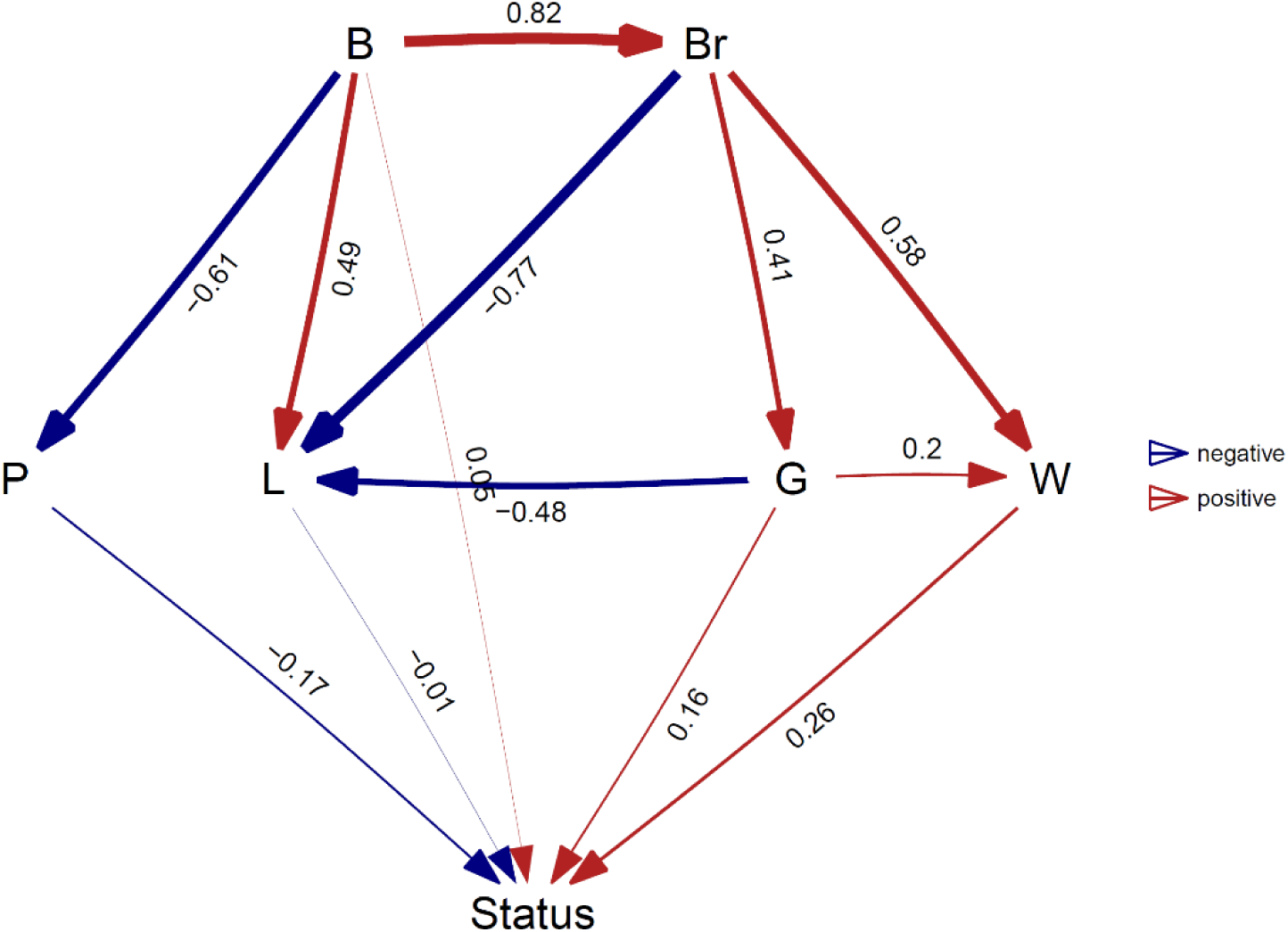
A visualization of the best supported causal model, and the standardized path coefficients.

A second way to look at a fitted model is to more directly look at the standardized coefficients and errors of the paths using `coef_plot`. We can use it to quickly compare the importance of the different variables that affect `Status`. Although we have modeled five effects on status, they are not necessarily all important and certainly litter size and body size have small effects (figure 4a).

**Figure 4:**
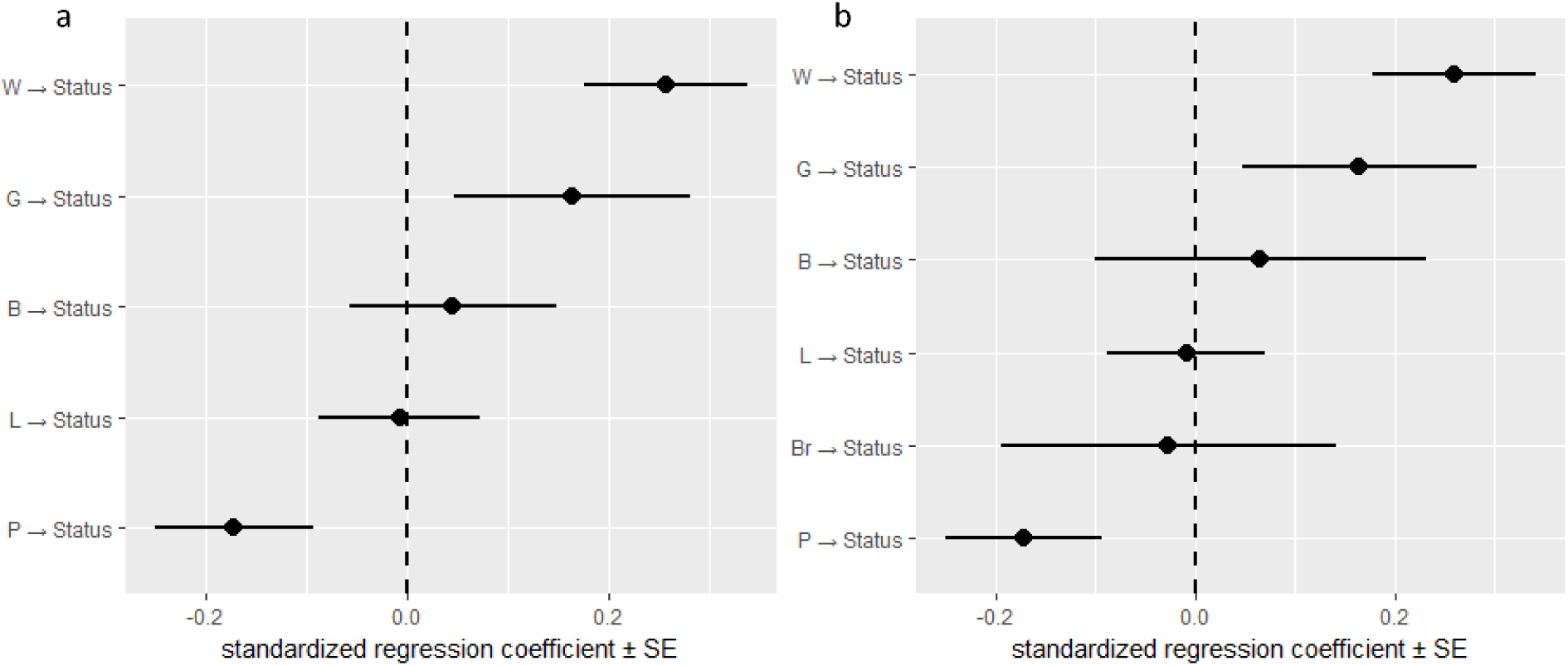
Standardized path coefficients and their standard errors, for the best supported model (*a*) and the average of the top two models (*b*).

~~~
**coef_plot**(b, error_bar = "se", order_by = "strength", to = "Status") + ggplot2::**coord_flip**()
~~~

### Model averaging

In many cases it may not be obvious or correct to choose one model. While in this case the two top competing models were nested, they do not have to be. In cases like these, it may be useful to perform model averaging instead, as discussed and used in the original paper (von Hardenberg & Gonzalez-Voyer 2013). `phylopath` makes model averaging easy, and you can quickly average over a selection of the top models, or all models considered. One should take care to not include models with significant C-statistics in the averaging, as these models are not supported. Models are weighted by their likelihood, and these weights can be found in the original `summary` table in the `w` column. One needs to choose how to deal with paths that do not occur in all models. One can average path coefficients only between models those include that path, this is often called conditional averaging and was used by von Hardenberg & Gonzalez-Voyer (2013) and is the default behavior in `phylopath`. Alternatively, one can consider missing paths to have a coefficient of zero and average over all models, which is often called full averaging. The latter results in *shrinkage*, where the path coefficients that do not occur in all models will shrink towards zero.

In this case, we could choose to average the two competing models. Let us use full averaging, and reevaluate the strength of the coefficients towards `Status` (figure 4b):

~~~
avg <- average(p, avg_method = "full")
coef_plot(avg, error_bar = "se", order_by = "strength", to = "Status") + ggplot2::coord_flip()
~~~

The average function selects the competing models, estimates the standardized path coefficients and then averages them. Note that we have only averaged over the two top models, since by default the cut_off is set to 2 CICc. You can average over all models in the set by using cut_off = Inf (but should only do so when all C-statistics are non-significant, see above).

### Analysis conclusion

A clear rejection of the null model indicates that brain size is related to the vulnerability to extinction of mammals, where large-brained animals high a higher vulnerability. This effect is mediated through life history, where the weaning and gestation periods are more important than litter size. There is no strong evidence in support of a direct effect of brain size on vulnerability to extinction that is independent of life history. The original analysis came to the same conclusion.

## Models of evolution

`phylopath` uses the `phylolm` package in the background (see below) and the models of evolution that are available there are therefore supported. You can simply pass the name of the model of evolution through the `model` parameter, just like using `phylolm` directly. It should be noted though, that `phylopath` by default uses Pagel's lambda model and not Brownian motion, which is the default for `phylolm`. Also, the model of evolution is only applied to continuous variables, i.e. using `phylolm::phylolm`, and not to binary variables which use `phylolm::phyloglm`. For the latter, one can choose between the two computational implementations, using the `method` parameter. When you supply the `model` or `method` parameter (or any other modelling parameters through the ellipses: …) to `phylo_path`, these settings are automatically passed down to other functions, `so best, choice` and `average` all use the same settings to guarantee consistency.

The estimated phylogenetic parameter can be found in the `d_sep` tables returned by `phylo_path` in the `phylo_par` column (we can also see which independence statements are rejected by looking at the p-values). For example, we can see the estimates of *lambda* for the `null` model above:

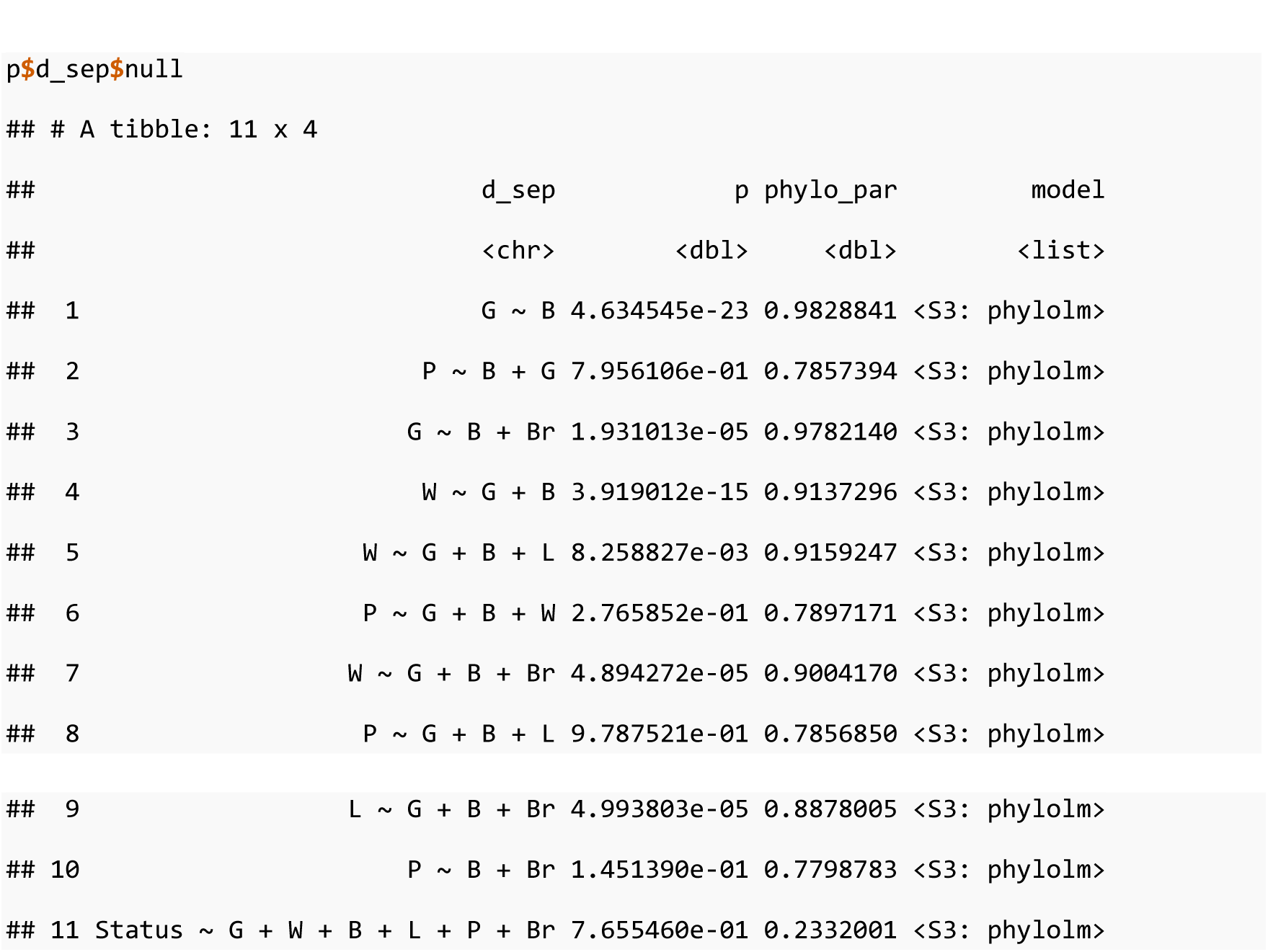

## Implementation notes

In addition to the functions outlined above, several lower level functions are also available to the user, specifically `est_DAG` to estimate the path coefficients of an arbitrary model, and `average_DAGs` to average several fitted models.

`phylopath` builds on several important packages, a few of which I highlight here. Firstly, it implements PGLS and phylogenetic GLM using `phylolm` (Ho and Ané 2014). This implementation was chosen for several reasons, including that the package is fast on large trees (an effect that is compounded in path analysis), its support for both Gaussian and logistic models, the robust estimation of confidence intervals using bootstrapping and the implementation of many standard S3 methods which makes it easily extendable by others.

Furthermore, the `ggm` (Marchetti et al. 2015) package is used for the ordering of the causal graphs and the finding of the d-separation statements. Model averaging is implemented using the `MuMIn` (Barton 2016) package. `ape` (Paradis et al. 2004) is used for checking and pruning phylogenies. `ggplot2` (Wickham 2016) and its `ggraph` (Pedersen 2017) extension are used for all plotting methods.

`phylo_path` supports parallel processing for analyses on very large phylogenies.

## Conclusion

I have presented `phylopath`, a package that aims to make phylogenetic path analysis more reproducible and less error-prone, and much faster and easier for the analyst. I hope that the package will stimulate the use of PPA amongst evolutionary biologists, as I believe that it is a powerful tool for a field in which experimental data is often impossible to obtain. I welcome bug reports, feedback and suggestions for the development of `phylopath`.

## Acknowledgements

I thank Alejandro Gonzalez-Voyer and Achaz von Hardenberg for their help during the development of the package. I thank Niclas Kolm and Alejandro Gonzalez-Voyer for their helpful comments on the manuscript and their support. `phylopath` would not exist without the R packages on which it depends or R itself, and the generous time and effort of those creating and maintaining open-source software.

